# Metaproteomics as a tool for studying the protein landscape of human-gut bacterial species

**DOI:** 10.1101/2021.09.02.458484

**Authors:** Moses Stamboulian, Jamie Canderan, Yuzhen Ye

## Abstract

Host-microbiome interactions and the microbial community have broad impact in human health and diseases. Most microbiome based studies are performed at the genome level based on next-generation sequencing techniques, but metaproteomics is emerging as a powerful technique to study microbiome functional activity by characterizing the complex and dynamic composition of microbial proteins. We conducted a large-scale survey of human gut microbiome metaproteomic data to identify generalist species that are ubiquitously expressed across all samples and specialists that are highly expressed in a small subset of samples associated with a certain phenotype. We were able to utilize the metaproteomic mass spectrometry data to reveal the protein landscapes of these species, which enables the characterization of the expression levels of proteins of different functions and underlying regulatory mechanisms, such as operons. Finally, we were able to recover a large number of open reading frames (ORFs) with spectral support, which were missed by *de novo* protein-coding gene predictors. We showed that a majority of the rescued ORFs overlapped with *de novo* predicted proteincoding genes, but on opposite strands or on different frames. Together, these demonstrate applications of metaproteomics for the characterization of important gut bacterial species. Results are available for public access at https://omics.informatics.indiana.edu/GutBac.

**Author summary:** Many reference genomes for studying human gut microbiome are available, but knowledge about how microbial organisms work is limited. Identification of proteins at individual species or community level provides direct insight into the functionality of microbial organisms. By analyzing more than a thousand metaproteomics datasets, we examined protein landscapes of more than two thousands of microbial species that may be important to human health and diseases. This work demonstrated new applications of metaproteomic datasets for studying individual genomes. We made the analysis results available through the GutBac website, which we believe will become a resource for studying microbial species important for human health and diseases.

## Introduction

It is now well established that microbial species inhabit many ecologies, which drives diversity due to the need to adapt to these environments^1^. Molecular diversity and robustness also allows these species to inhabit microbiomes that are host associated, such as the human gut^2–7^. Due to its importance to human health and disease, the human gut microbiome has been extensively sequenced, leading to the identification of more than a thousand distinct species, with knowledge about diversity increased by every new study^8^. Culture based whole genome sequencing techniques^9,10^ have identified a few hundred human gut associated genomes. Culture free techniques, revolutionized by metagenomics coupled with computational genome binning methods, have resulted in many more metagenome-assembled genomes (MAGs)^1,3,5,8–11^. A unified genome catalog contains more than 200,000 reference genomes from the human gut microbiome^8^. Maintaining a comprehensive genomic catalog of bacteria and archaea will provide the basis needed to perform large scale multi-omic comparative genomic studies. Large scale studies based on whole genome sequences will be central in understanding details about mechanisms of microbial interactions with each other and with their environments and hosts. These studies will also be critical for uncovering details about metabolic pathways and key functions at the protein level by examining proteome landscapes and employing various data-mining techniques to identify genes and functions of interest, such as CRISPR-cas systems^12^ and anti-CRISPRs^13^.

Recent progress has dramatically increased the collection of microbial species that are related to human health and diseases, most notably the accumulation of MAGs, many of which represent new species. Experimental studies of these new species in terms of their expression and functions remains scarce. Computational gene predictors have become an essential first step in the annotation of these new genomes. *De novo* gene prediction techniques are commonly used because they are not constrained by sequence similarity with known ones^14,15^. *De novo* gene prediction remains a unsolved problem, with different tools, such as FragGeneScan, prokka, and GenMark, producing largely consistent but not perfect predictions because most of the predicted genes remain hypothetical without functional annotations. Proteomic studies have been used to improve understanding of the microbial world beyond genomics. Proteomics allows correction of bad gene predictions, and the discovery of protein products from the regions of the genome not yet predicted to be coding areas^16^. Proteomics has been used a tool for studying bacterial virulence and antimicrobial resistance^17^. The Multidimensional Protein Identification Technology (MudPIT) approach was used to study *Pseudomonas aeruginosa* membrane-associated proteins, which contribute to *P. aeruginosa* cells’ antibiotic resistance and is involved in their interaction with host cells^18^.

Motivated by recent expansion of microbial genome catalogues, our previously defined reference based peptide identification pipeline (HAPiID)^19^, and the increasing number of publicly available gut metaproteomics datasets, we conducted a large scale survey of metaproteomics data of the human gut microbiome to study the proteome landscapes of the various microbial species dominating the human gut. Our aim was to mimic targeted proteomics studies, which traditionally focused on single cultured species, by leveraging the available metaproteomics datasets. We were able to identify species that are ubiquitously expressed across all samples spanning various phenotypes. Furthermore, by focusing on the most highly expressed genome sequences at the protein level, we represented the expressed proteome as a network and extracted co-abundant protein modules. We used such network information to study the various metabolic pathways that a protein with unknown function might be involved in and also identified modules that were expressed in hosts with specific phenotypes. We also leveraged proteome information to identify and annotate potential operon structures within genomes and recover open reading frames (ORFs) with spectral evidence that were otherwise missed by computational protein coding gene predictors. Operon structures have been exploited for computational functional predictions (guilt by association)^20,21^. We made the results from our analysis available at a publicly accessible website, providing the protein expression and putative operon structures with spectral support for many human gut related microbial species for the first time.

## Results

### Not all highly-expressed species are equal: some are generalist and some are specialist

Taking advantage of the large number of metaproteomics datasets, we were able to identify a total of 2,511 distinct genomes that were expressed in at least one out of the total 1,276 samples (see Table 1). A total of 13,460,264 spectra were matched to peptides, out of which 12,950,155 spectra (96.2%) were matched to the 2,511 highly abundant genomes. The remaining 510,109 identified spectra were not considered further in this study. Figure 1 summarizes the spectral support for these genomes with x-axis showing the number of supporting samples and y-axis showing the total number of identified spectra for each genome (a data point in the plot).

**Table 1:**
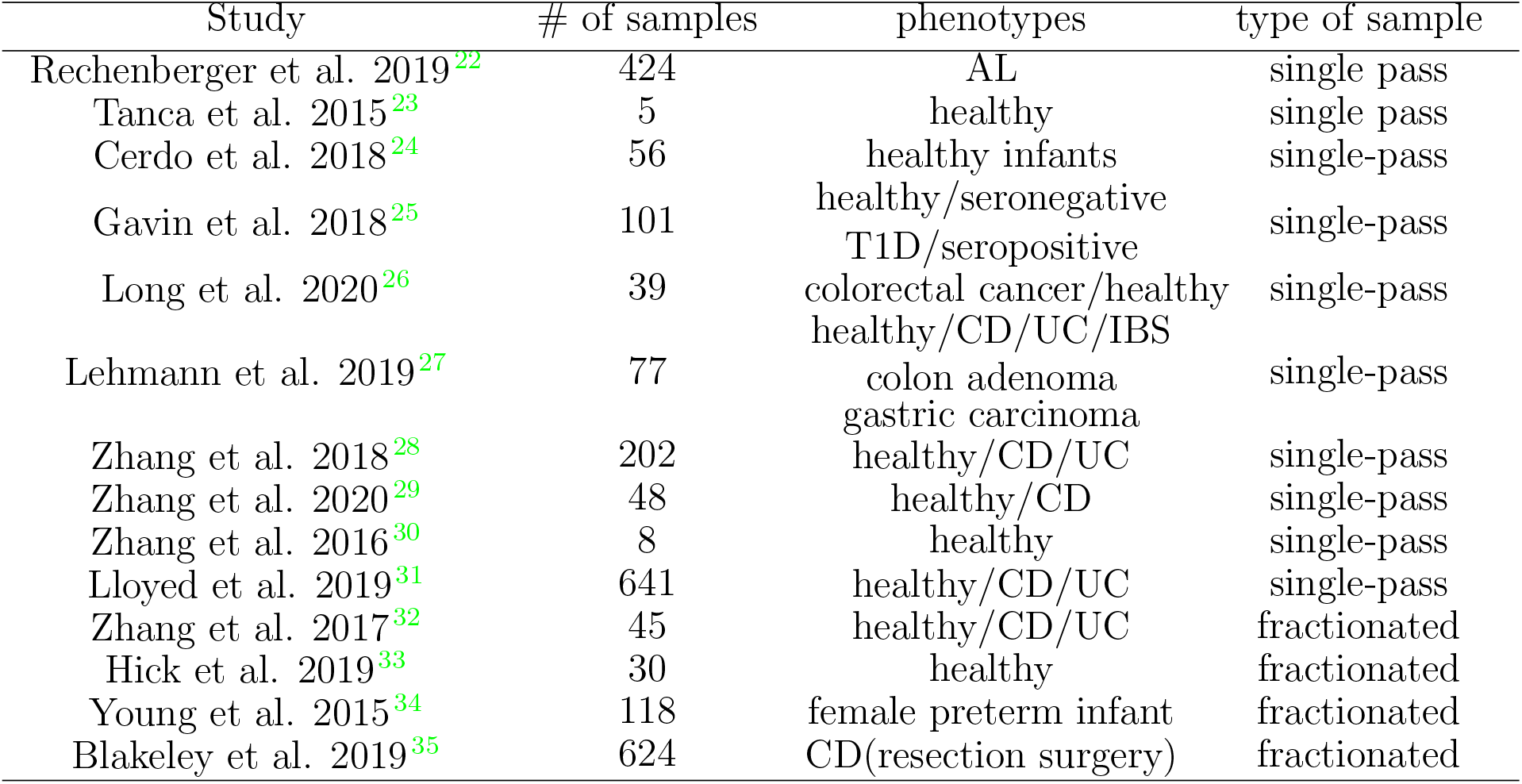
Summary of the metaproteomics datasets that were analyzed.

**Figure 1:**
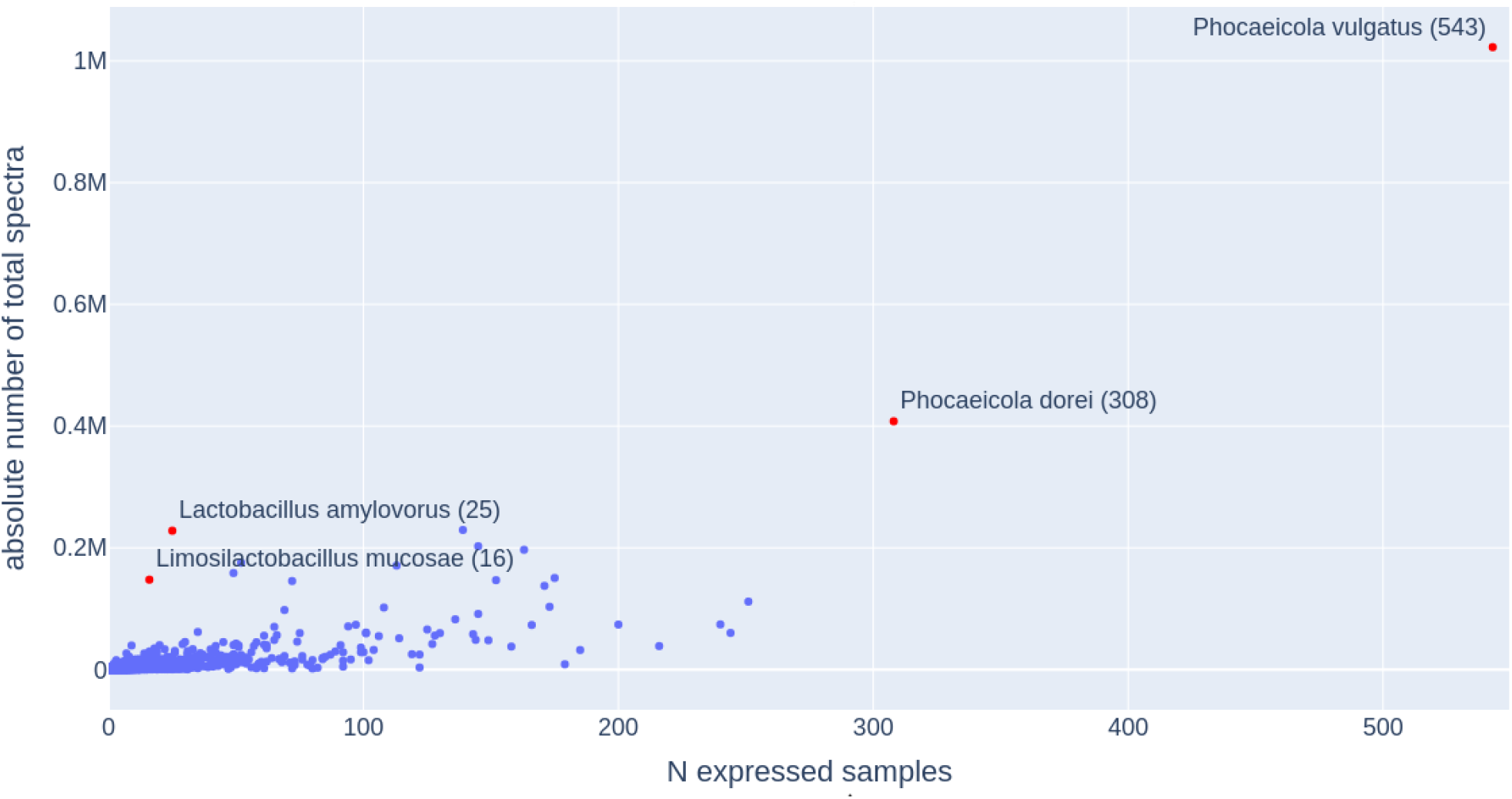
Scatter plot summarizing the expression of human associated microbial species in different samples. X-axis shows the number of samples in which each genome was observed, and y-axis shows the total support spectra for each genome. The four genomes that are highly expressed and/or broadly distributed are highlighted in red with their species names shown in the plot, followed by the number of samples supporting their expression in the parenthesis.

We saw variation in metaproteomic support for the different genomes with the most highly expressed genome having over one million supporting spectra. The average number of spectra expressed by each genome was 4,780 and the mean was 385 spectra per genome, which indicates very few highly expressed genomes and many low expression genomes at the protein level. To improve coverage at the protein level, we focused on the top 100 most abundant genomes for the rest of this section. The top 100 most abundant genomes (less than 4% of the total number of expressed genomes) contributed more than 61% of the identified spectra (7,327,171 spectra). Figure 2 summarizes the taxonomic composition at the genus level of these top 100 genomes. These 100 genomes represent a total of 34 genera in five most abundant phyla of the human gut *Bacteriodetes, Firmicutes, Actinobacteria, Proteobacteria* and *Verrucomicrobiota* (Supplementary Figure S1).

**Figure 2:**
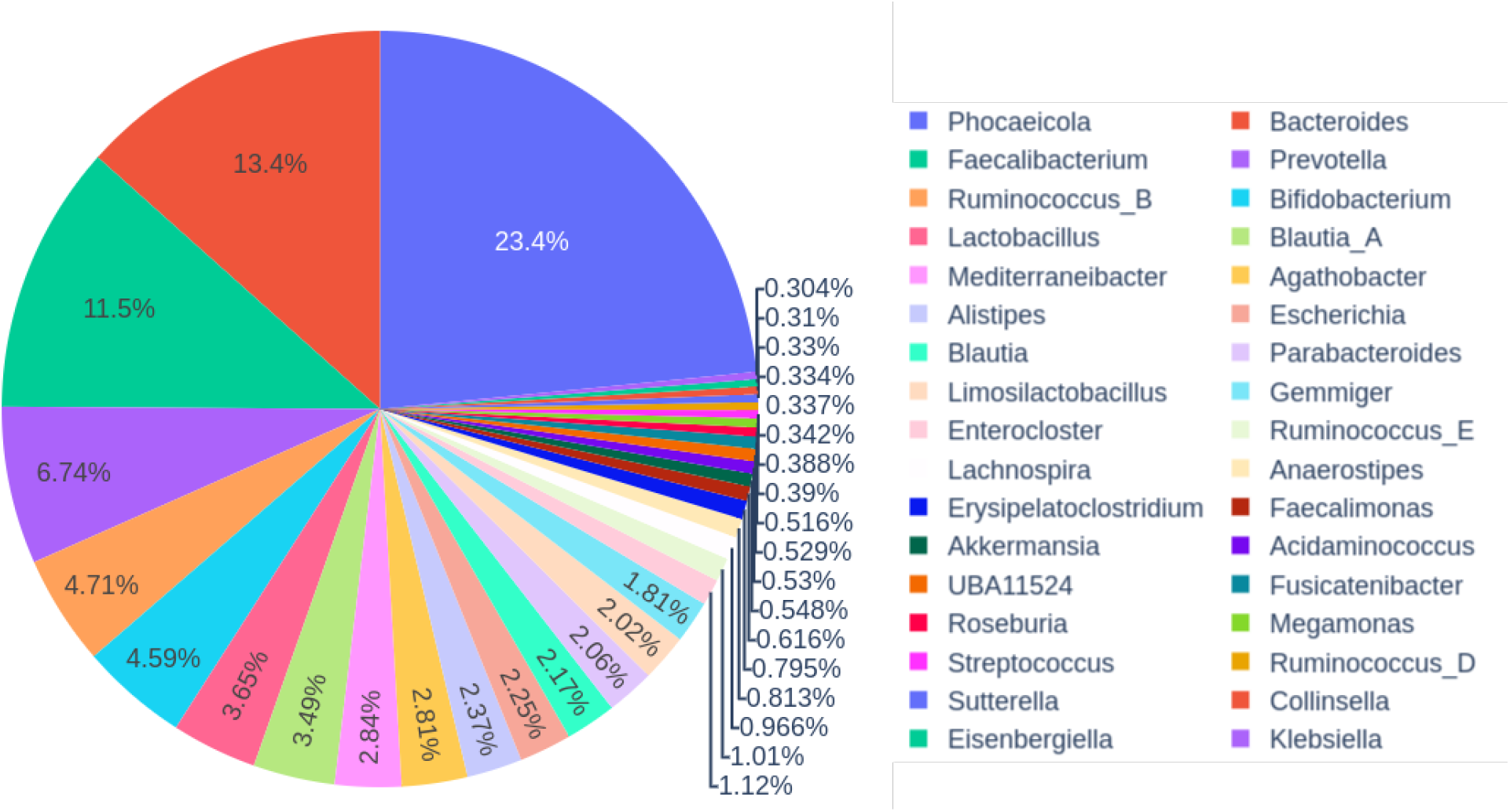
Piechart summarizing the taxonomic composition at the genus level for the top 100 highly expressed genomes.

The top two most abundant species belong to the genus *Phocaeicola* (*Phocaeicola vulgatus* and *Phocaeicola dorei*, which expressed more than 11.9% of the total number of spectra in a total of 543 and 308 samples respectively. We refer to these two species as generalists herein forward. These two species share similar phenotype expression patterns because they both appear in many healthy samples and in similar disease samples, including AL, T1D, CD, CD, IBS, colon adenoma and CD (followed by resection surgery), with the exception of gastric carcinoma where only *Phocaeicola vulgatus* was found to be expressed. We also identified two species belonging to the *Lactobacillaceae* genus (*Lactobacillus amylovorus* and *Limosilactobacillus mucosae*), which were the 4^*th*^ and the 11^*th*^ most highly expressed genomes but are only expressed in 25 and 16 samples respectively. We refer to these two species as specialists. The latter two species were not found to be expressed in a single healthy sample, but both were highly expressed in CD (followed by resection surgery) patients, and the former was also found in AL patients with high abundance.

We next checked whether or not these two pairs of species have similar functional profiles when compared with each other, over the samples in which they are expressed. To do that, we annotated their expressed proteins, as much as possible, with COG terms (see methods for more details). For each of the four genomes mentioned above, we computed the relative abundances for the COG terms associated with their expressed protein sequences. We summarize the distribution of the high level single COG categories, represented by 25 single letter groups, in Figure 3. We noticed that, overall, the two generalist species share more similar functional profiles, and the two specialist genomes share more similar functional profiles (Figure 3). The two specialist genomes had significantly higher relative protein expression levels for the COG categories G (carbohydrate transfer and metabolism), J (translation, including ribosome structure and biogenesis) and T (signal transduction). On the other hand, the two generalist species had relatively higher expression levels in the COG categories P (inorganic ion transport and metabolism), M (cell wall/membrane/envelope biogeneis), C (energy production and conversion), U (intracellular trafficking, secretion, and vesicular transport) and W (extracellular structures). It should be noted that the latter functional category W had no observed expression within the two specialist species.

**Figure 3:**
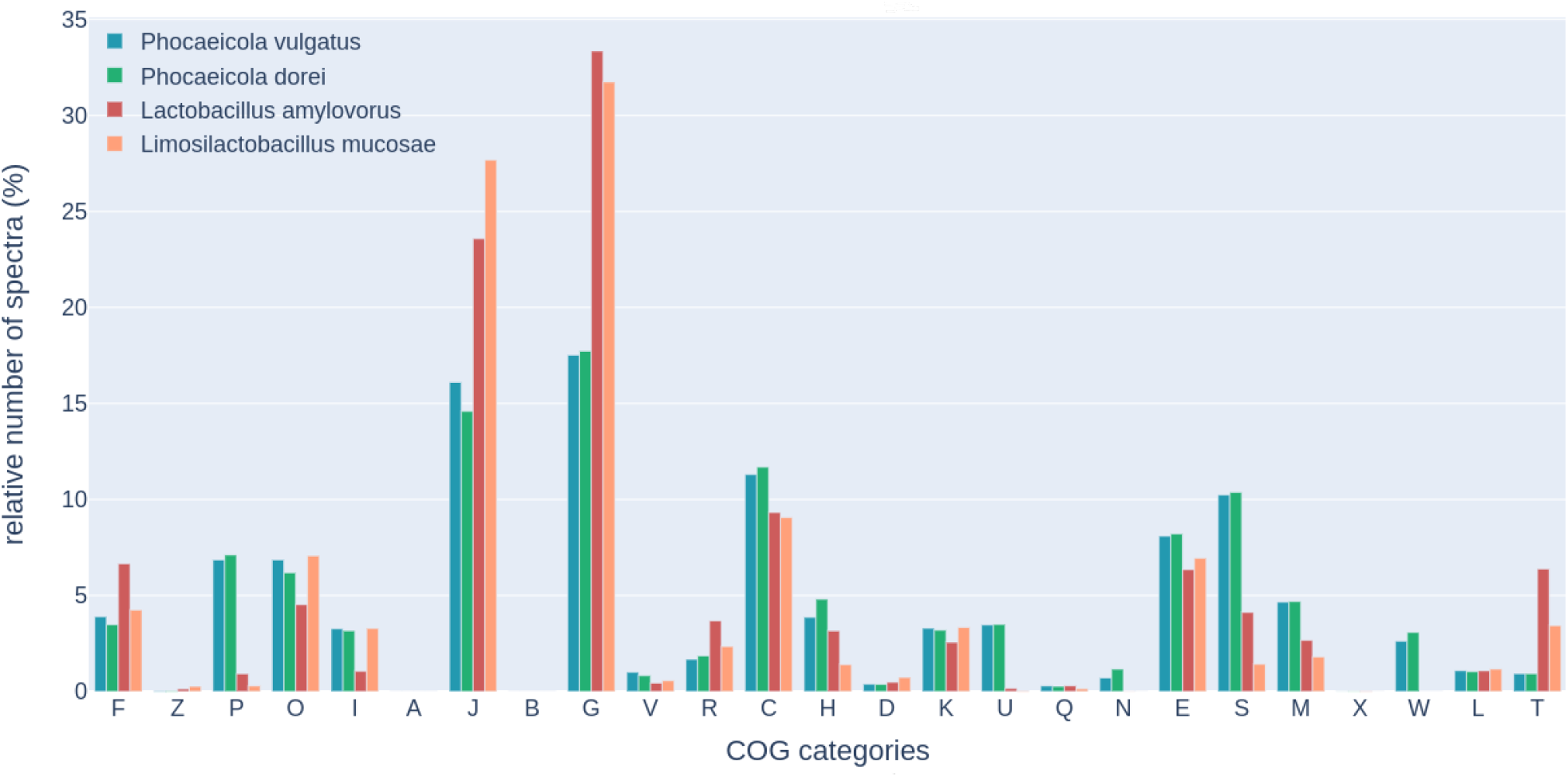
Barplot summarizing the relative abundances of the COG functional categories of the two generalist (blue/green) and two specialist (red/orange) highly abundant genomes

### Protein co-expression modules and their applications

We focused on the top 100 most highly expressed genomes in this section to assess the presence/expression of their proteins among the different host phenotypes (see Supplementary Figure 2 for the summary). We derived groups of proteins (protein co-expression modules) that had similar presence/expression patterns across samples and were mostly found in one of the 13 phenotypes. We extracted a total of 854 such modules, composed of 3,697 protein sequences (see Table 2). Figure 4 shows two such protein modules. The first module containing proteins that were mostly exclusively expressed in CD patients followed by resection surgery and the second one contains proteins mostly expressed in patients with acute leukemia. We noticed that many of the proteins within the modules lack functional annotations but have connections to those whose functions are previously characterized.

**Table 2:** Phenotype specific protein co-expression modules

| Phenotype | # of modules | # of proteins |
| --- | --- | --- |
| AL | 232 | 1,250 |
| CD | 96 | 307 |
| CD (resection surgery) | 225 | 816 |
| colon adenoma | 6 | 17 |
| gastric carcinoma | 9 | 23 |
| healthy | 169 | 873 |
| healthy infant | 25 | 92 |
| IBS | 6 | 26 |
| preterm infants | 0 | 0 |
| T1D | 12 | 45 |
| UC | 74 | 248 |

**Figure 4:**
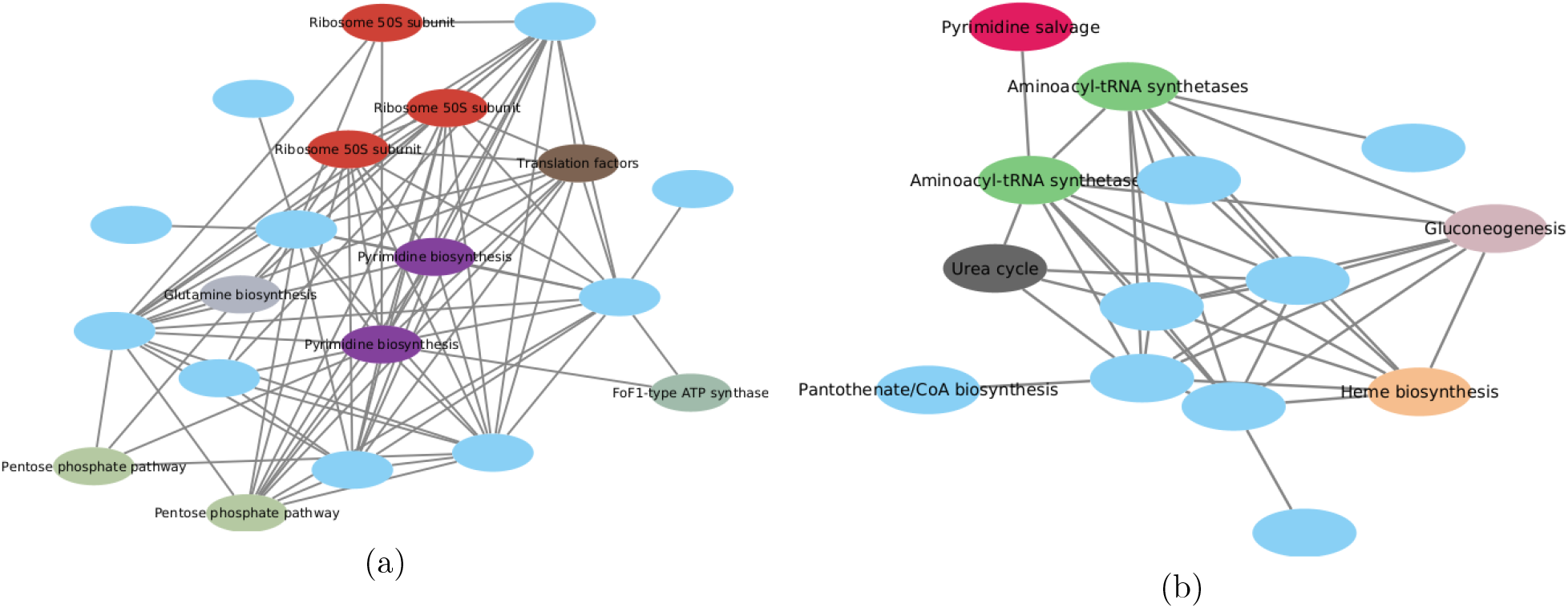
Protein co-abundant modules extracted from (a) *Bifidobacterium longum* that is mostly expressed in CD (followed by resection surgery) patients and (b) *Escherichia flexneri* mostly expressed in acute leukemia patients.

We explored the possibility of suggesting functions for proteins that lack COG or KEGG annotations, but are co-expressed with other annotated proteins (guilt-by-association). We report pathway associations when such proteins share edges with other annotated ones within the modules. By doing so we were able to suggest potential pathways for a total of 7,682 proteins using COG annotations and 5,800 proteins using KEGG annotations (9,566 receiving either COG or KEGG pathway). On the other hand, there were a total of 3,747 proteins using COG annotations and 2,080 proteins using KEGG annotations respectively, that either had no edges with any other annotated proteins or were found in modules that were completely composed of un-annotated protein sequences.

### Putative operon structures with spectral support

We applied the described pipeline to extract potential operon candidates from genomes that were expressed in relatively high number of samples (see Methods for more details). Two lists of potential operons were produced: one with spectral support (as described in the methods section), and the other one of operons without consideration of spectral information. In total, we analyzed 278 genomes, and made the results available on the GutBac website. From these genomes, a total of 4,089 potential operon structures with spectral support and 36,633 suggested operon structures with or without spectral support were identified. We compared our predictions with those predicted using fgenesB for the top 10 most highly expressed genomes, and the results are summarized in Table 3. The number of potential operons with spectral support were always lower than those predicted by fgeneB and those suggested by our pipeline without spectral support. However, on average there was slightly better agreement between the fgenesB predictions and those with spectral support (*>* 92%) compared to the suggestions without spectral support (88%). It should be noted that the increase in the difference between the reported numbers is expected in this case, as we included more genomes with decreasing spectral coverage.

**Table 3:**
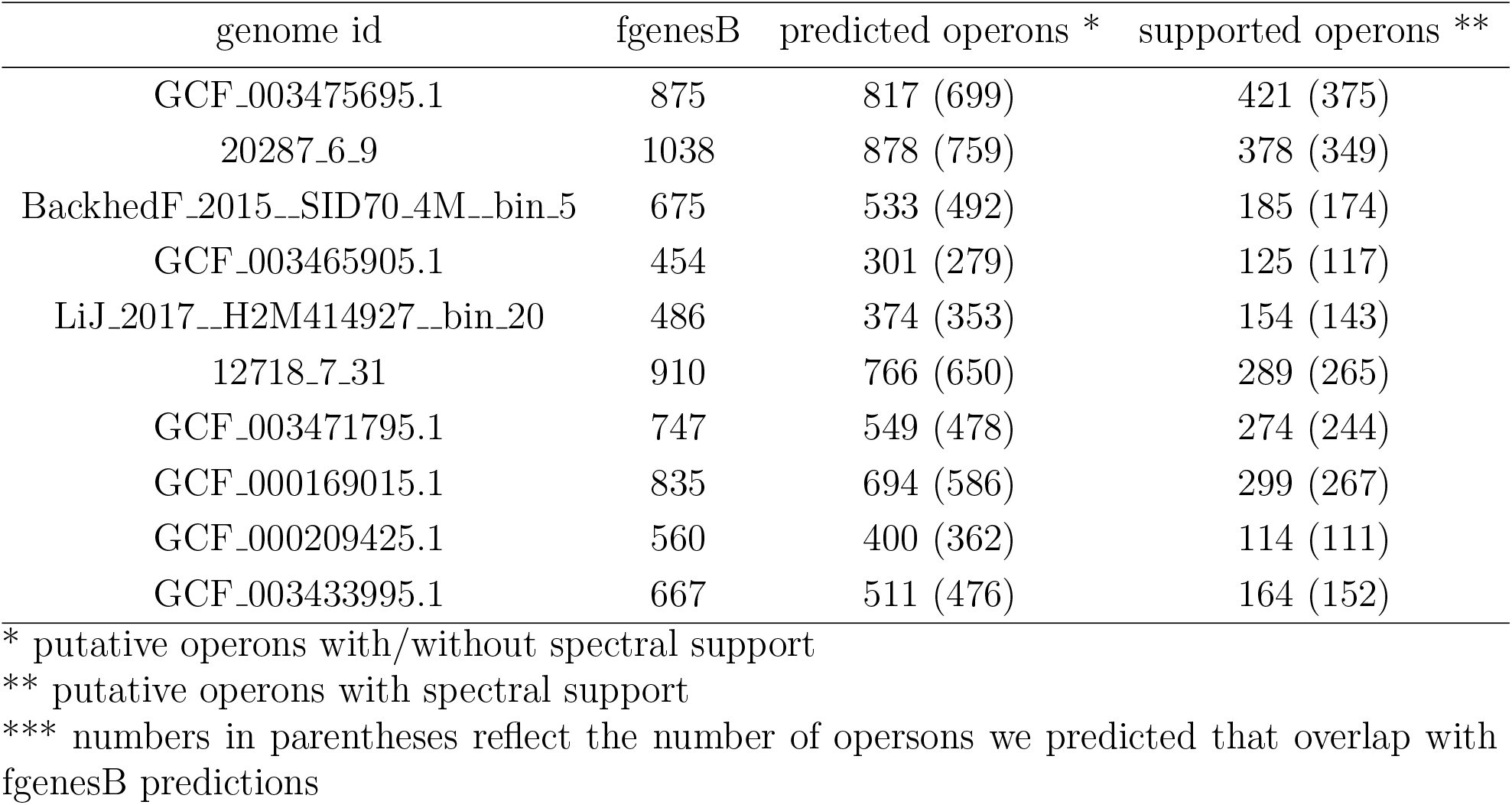
Summary of predicted operons by fgenesB and our approaches using spectral support and without spectral support for selected genomes.

### Recovery of missed ORFs with spectral support

Here we incorporated metaproteomics datasets to recover ORFs with spectral support which were otherwise missed by *de novo* gene predictors mentioned in the methods section above. Novel ORFs in comparison to the two computational gene/protein prediction methods (FragGeneScan and prokka) were extracted as explained in Methods. There were a total of 2,463 distinct genomes that were identified to be highly expressed at the protein level in at least 1 sample. A total of 23,949 putative ORFs were identified which were otherwise missed, when FGS was employed for gene prediction. Similarly, a total of 22,658 novel ORFs were recovered when prokka was used for gene prediction. The majority of these recovered ORFs were overlapping (at 84%); a more detailed results for this comparison is summarized in Supplementary Figure 3. Unsurprisingly, there is positive correlation between the number of ORFs recovered for genomes with their protein expression levels in both cases, as shown in Supplementary Figure 4. This further suggests potential improvement in bacterial annotations with the increase in the throughput of metaproteomics.

We examined the relationship of the rescued ORFs with respect to the protein coding genes predicted by FGS or prokka. We found that among the rescued ORFs (22,573), the majority of them are either on the opposite strands (11,955, 53%) of already predicted protein coding genes, or the same strand but of different frames (7,347, 33%). The genome that has the most number of rescued ORFs is *Phocaeicola vulgatus* (accession ID: GCF 003475695.1), one of the two generalists we discussed above. A total of 426 ORFs were recovered (comparing to prokka prediction), among which 232 cases were found on the opposite strands of genes predicted by prokka, and 97 were found to be encoded by a different frame of overlapping gene. Figure 5 shows plots of three regions in this genome that contain rescued ORFs with spectral support (together with predicted operons). The first example (illustrated on the top in Figure 5) involves two rescued ORFs, among which one is a large ORF that was missed by prokka (but was predicted as a protein coding gene by FragGeneScan), and could be re-identified using metaproteomic data (we note this ORF was missed by prokka probably because it overlaps with a tRNA gene, shown as a red arrow in the plot). The second example (shown in the middle in Figure 5) also contains two rescued ORFs, and we note the first ORF (from 86972 to 87128 bp in NP QRPW01000005.1) was missed by both FragGeneScan and prokka but we found metaproteomic evidence for this ORF. In fact, this ORF is part of a large operon that involves genes encoding ribosomal proteins, and hmmscan search (https://www.ebi.ac.uk/Tools/hmmer/search/hmmscan) shows that the rescued ORF is ribosomal protein S11. In the last example, the larger rescued ORF of 199aa overlaps (on the opposite strand) with two putative protein coding genes. This ORF is supported by one peptide of 2 PSMs but similarity search (using hmmscan search) did not return similar sequences, so further investigation could be needed.

**Figure 5:**
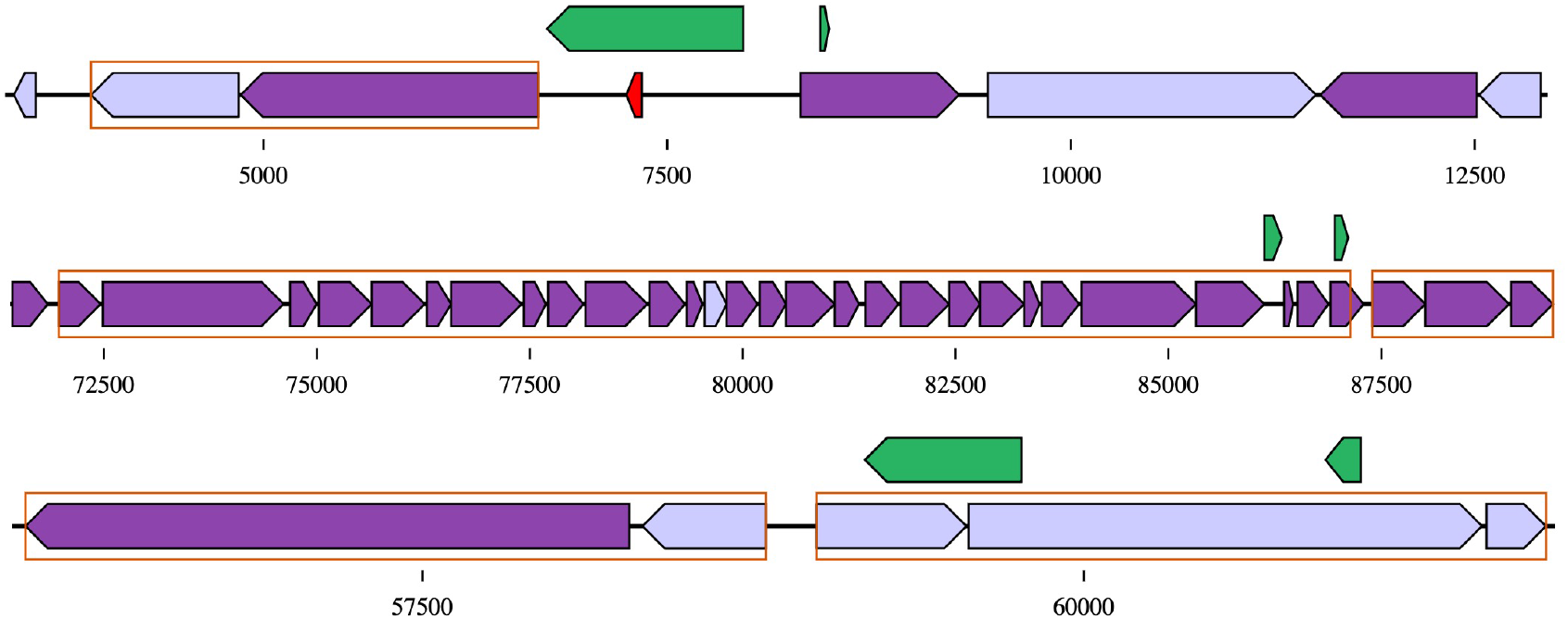
Selected cases of rescued ORFs using metaproteomic data in *P. vulgatus* genome. The three blocks of arrows represent genes predicted from three regions in this genome: from the top to the bottom are contigs with IDs of NZ QRPW01000017.1, NZ QRPW01000005.1, and NZ QRPW01000002.1, respectively. Genes predicted by FGS and/or prokka are shown as arrows in dark purple and light purple for genes with spectral support in at least two metaproteomic datasets, or only one metaproteomic dataset, respectively (the red small arrow represents a tRNA gene predicted by prokka) around the central lines, each representing a segment of the genome. Rescued ORFs are shown as green arrows above the lines. Genes in the same putative operon structure are surrounded in orange squares.

## Discussions

By taking advantage of the availability of many metaproteomics datasets, we were able to probe the protein landscapes for many human-associated microbial species. However, due to the relatively low throughput of metaproteomics comparing to metagenomics (and metatranscriptomics), the number of genomes we studied (in depth) was rather limited, even though a huge number of reference genomes for human gut microbiome were available. The HAPiID pipeline includes more than 6000 genomes for peptide and protein identification from metaproteomic data, and we were only able to provide genome-level protein landscape analysis for 40% of these genomes.

Using metaproteomic data, we were able to identify a large number of ORFs that had spectral support but were missed by *de novo* gene predictors. Many of these rescued ORFs are relatively short (otherwise they are unlikely to be missed by protein coding gene predictors), and we want to emphasize that they need to be interpreted cautiously. Some of them could be false identifications and some of them may reflect translation that does not result in proteins with biological significance. In the case of the 199aa ORF discussed in the results, evidence of this ORF was based on a single peptide of 2 PSMs and more focused mass spectrometry analysis (higher throughput techniques or a focus on this species) could be used to improve support for this existence of this ORF and eliminate the possibility of it being a false identification. Finally, we hope the finding of “rescued” ORFs (such as the one that overlaps with a tRNA gene) and their analysis can inspire ideas for improving *de novo* protein coding gene predictions.

With the increasing number of microbial genomes being sequenced, functional annotation becomes an immediate need. Numerous genomes are being computationally assembled as a result of these metagenomic sequencing efforts coupled with emerging computational genome assembly and binning tools. This is expanding the gap between the amount of whole microbial genomes recovered and the fraction of annotation the community has concerning these newly discovered genomes. While metaproteomics based techniques are still low throughput compared to the sequencing base techniques, we believe there is great value in studying the proteome of these microbial communities directly within their environment and augmenting a third level of information on top of metagenome and metatranscriptome, to capture such microbial interactions at a more granular scale, i.e., both functional and pathway levels, rather than a mere interpretation of the genome abundance levels of the different microbial entities at some taxonomic level.

We hope that our results would serve as a resource for the study of gut microbial community in particular to cast more light over the microbial dark matter, and broaden our understanding at the functional level. Furthermore, we also hope that this work would inspire others and serve as an example on how to utilize metaproteomics as a tool for large scale analysis to study the microbial functional landscapes, especially as metaproteomics throughput improves and different mass spectrometry methods that may improve results, such as multiplexing and data independent approaches, become more common.

## Materials and Methods

### Metaproteomics samples

Human gut metaproteomic datasets were obtained from the publicly accessible proteome exchange database^36^. We extracted a total of 2,418 Thermo Fisher RAW files, from 14 recent studies, spanning 12 distinct phenoypes^22–35^. A more detailed summary of the datasets used can be found in Table 1. Four of the studies were based on fractionation approaches to increase sequencing depth. For peptide-spectral-matching and to identify expressed proteins, we ran these individual samples through our previously developed HAPiID pipeline^19^. For fractionated samples, we performed peptide spectral matching for each individual fraction separately and then combined the identification results from every fraction into results for a single sample.

### Peptide identification and proteome quantification

To identify peptides and quantify proteome content of our proteomics dataset, we first ran each individual sample through our previously implemented HAPiID proteomics framework (https://github.com/mgtools/HAPiID)^19^. Default parameters were used over all samples. For the peptide spectral matching step, the MSGF+ search engine was used with the following settings^37^: high-resolution LTQ as the instrument type, precursor mass tolerance of 15 PPM, isotope error range between -1 and 2, a maximum of 3 fixed modifications, variable oxidation of methionine, fixed carboamidomethyl of cysteine, maximum charge of 7 and a minimum charge of 1, and allowing for semi-tryptic fragmentation up to two missed-cleavages. A protein database from proteomes of 6,160 non-redundant microbial whole genome sequences was used as a reference to compute theoretical spectra, for peptide spectral matching. The microbial genomes were collected from five recent studies^3,5,9–11^, and then filtered and dereplicated by dRep^38^, using 90% sequence identity for the primary clusters, and 99% sequences identity for the secondary clusters with a minimum of 60% genome alignment coverage. A strict FDR cutoff at 1% was enforced by using target-decoy database approach with reverse protein sequences as decoy. A final set of samples was maintained by discarding the ones where we identify less than 1,000 unique peptide sequences. After combining fractions for the fractionated samples, our final dataset was composed of 1,276 samples.

Highly abundant genomes at the protein level for each sample was defined as the list of top *N* genomes that were able to cover 80% of the identified spectra in the first step based on the greedy approach defined in the HAPiID pipeline^19^ (see^19^ for more details). After selecting a list of highly expressed genomes for each sample, identified spectra gets redistributed across these genomes. Initially, all uniquely mapped spectra get assigned to their corresponding genomes, followed by partial allocations of the multi-mapped (i.e. shared) spectra. Quantities from unique spectra to genomes are used for weighted assignment of the multi-mapped spectra shared between multiple genomes, similar to the approach proposed in Qin et al.^39^. Genome to absolute spectral counts were later normalized by the respective proteome sizes (calculated as the total combined lengths of protein sequences for each genome), and then normalized by the total number of spectra for each sample to account for sequencing depths and allow for cross sample comparisons.

### Protein function annotation

We used both COG^40^ and KEGG^41^ databases to assign functional annotations to protein sequences. The latest version of clusters of orthologs (COG) database (release 2020), was used to assign COG terms, with significant hits, to the predicted protein sequences from whole genome sequences. A COG protein reference database was first created from the curated list of 3,213,025 protein sequences with COG assignments. Diamond blast^42^, with the *–more-sensitive* setting, was employed to assign COG terms to our query sequences. A protein sequence was annotated with a COG term if the alignment between the query protein sequence and the reference protein sequence covers at least 50% of the total length of the COG domain, with a minimum e-value score of 0.01. A total of 10,546,105 protein sequences (out of 15,072,008 protein sequences), each had at least one COG hit, using these criteria. Protein sequences that did not receive any COG hit were assigned to the category *S* (i.e., no COG-term assignments). We also performed COG based pathway annotation, for protein sequences with COG hits that participate in certain pathways. To transfer KEGG annotations to our protein sequences we first annotated protein sequences with KO terms using KEGG’s blast-Koala tool, over the latest version of KEGG database^41^, followed by mapping putative pathways that each protein participates in using KEGG’s pathway mapper tool. Furthermore, we post-processed protein to KEGG pathway maps by using our previously developed tool, MinPath^43^, to overcome potential overestimation of pathways by the naive function-to-pathway mapping method.

### Inference of protein co-expression network

For the top 100 most highly expressed genomes, we created protein expression profile *M* x *N* matrices where M represents the list of expressed proteins and N represents the list of samples where the particular genome is expressed in, and a particular entry *m*_*i*_, *n*_*j*_ represents the number of spectra expressed by the protein *m*_*i*_ in the sample *n*_*j*_. From this matrix, protein expression profile vectors were then extracted for each expressed protein and all against all pairwise correlation coefficients were calculated using the program fastspar^44^, which is a fast and scalable implementation of the original sparCC correlation measure^45^. The resulting *M* x *M* correlation matrix is used to construct a protein co-abudnance network. Based on author recommendations, we used a minimum sparCC correlation coefficient of 0.3 or higher to infer an edge between two expressed proteins. To extract proteome connected components, we developed an in house script using python’s networkX module^46^. A graph node in this context is an expressed protein sequence, and an edge between two protein sequences represents a sparCC correlation coefficient of 0.3 or higher, between the expression profile of these two proteins. We first identified all the maximum cliques within the network by an iterative approach. For all the remaining nodes that did not form cliques, we identified the best candidate connected components, by adding them to the clique where they share the highest number of edges with. The latest version of cytoscape was used for network visualization^47^.

### Host phenotype specificity of gut bacterial proteins

To quantify if a protein’s expression is only observed in gut microbiomes with a specific host phenotype or broadly found in samples with different phenotypes, we defined the host phenotype specificity (HPS) of a protein using Shannon’s entropy measure^48^, as following:

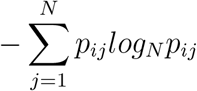

where *N* is the total number of possible phenotypes that a protein *i* was found to be expressed, and *p*_*ij*_ represents the proportion of the protein *i* being expressed in sample with phenotype *j*. The proportions for each protein to be found expressed in a phenotype is calculated based on the spectral counts, normalized by sequencing depth, for a protein across the the samples with different phenotypes. Using this metric, proteins with phenotypic entropy closer to zero indicates phenotype specific expression patterns, and those with values closer to 1 indicate more broadly distributed protein expressions, likely to be expected in a broad range of phenotypes.

### Inference of operon structures with protein expression support

Protein spectral support was integrated with genomic context to extract potential candidates for operon structures within highly expressed genomes. For each genome, we extracted clusters of proteins from the same contig and strand containing member protein sequences within 100 bases. We integrated spectral support to filter out clusters that did not have at least half of its protein members expressed in 2 or more samples and required at least 2 spectra per sample. To validate our results, we also predicted operon structures using the fgenesB program^49,50^. FgenesB is a bacterial operon and gene prediction program, based on pattern Markov chains. The FgenesB software suite, however, only provides a web interface with limited submissions per day per user, therefore an automated method to detect operon structures, that could be integrated in pipelines and run offline would be highly desirable. For comparison, we counted a predicted operon as overlapping with those of fgenesB if the two suggested operon regions on the genome overlap by more than 70%.

### Recovering of proteins missed by gene predictors but are supported by metaproteomics data

FragGeneScan (FGS)^51^ was used to predict protein coding genes for the genomes used in our reference protein database. For each of the highly abundant genomes identified in each sample, by selecting top N genome sequences covering 80% of the identified spectra in step 1 of our HAPiID pipeline (see^19^ for more details), we construct a protein database using 6 frame genome translation sequences instead of predicted protein sequences as a reference database. We then identified peptides from each sample based on this method of database construction. For each new peptide we extracted open reading frames (ORFs) that surround them, and then filtered out those that did not have a blast hit of 70% or higher with the respective predicted protein predicted by FragGeneScan. Similarly, we repeated the same task by using prokka^52^ for protein prediction instead of FragGeneScan.

### Availability of the results

Results are available through the GutBac website at https://omics.informatics.indiana.edu/GutBac/. The website includes GFF files with predicted proteins for each genome, FNA and FFN files containing relevant genomic sequences, GFF files containing both the predicted proteins and the missed ORFs for each genome, and CSV files containing the list of predicted operons for each genome. The website includes contig-specific plots generated by DNA Features Viewer^53^ containing the missed ORFs, predicted proteins, and predicted operons for both FGS and prokka. The website also features genome-specific searching and filtering.

## Supporting Information Legends

**Figure S1:** Piechart summarizing the taxonomic composition at the genus level for the top 100 highly expressed genomes.

**Figure S2:** Boxplots summarizing the host phenotype specificity of expressed proteins encoded by the top 100 most abundant genomes. Gray line indicates the average host phenotype specificity of the proteins in each genome.

**Figure S3:** Venn diagram summarizing the overlap between the rescued ORFs that were missed by FragGeneScan and prokka.

**Figure S4:** Scatter plots showing the relationship between the expression levels of the different genomes at the protein level with the total number of rescued ORFs. (A) rescued ORFs missed by FragGeneScan; (B) rescued ORFs missed by prokka.

