## Supplementary Tables and Figures for "Metaproteomics as a tool for studying the protein landscape of human-gut bacterial species"

### Supporting Information

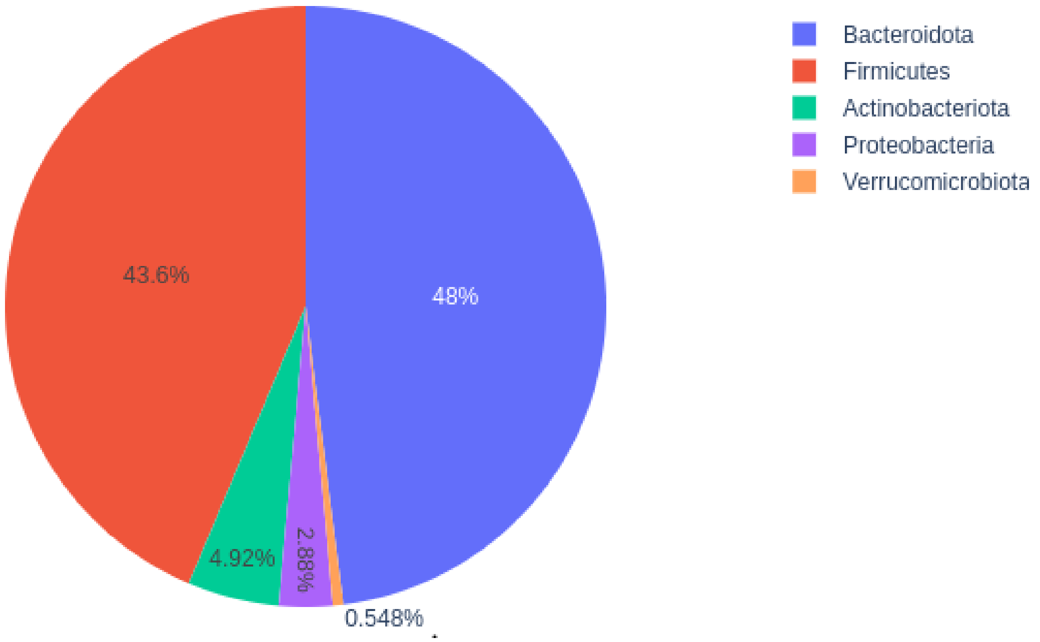


Figure S1: Piechart summarizing the taxonomic composition at the genus level for the top 100 highly expressed genomes.

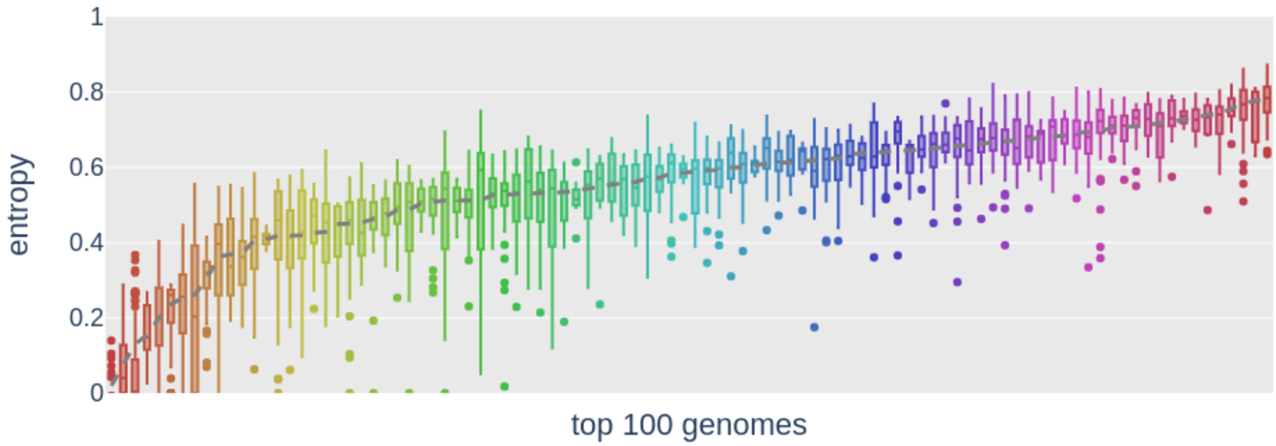


Figure S2: Boxplots summarizing the host phenotype specificity of expressed proteins encoded by the top 100 most abundant genomes. Gray line indicates the average host phenotype specificity of the proteins in each genome.

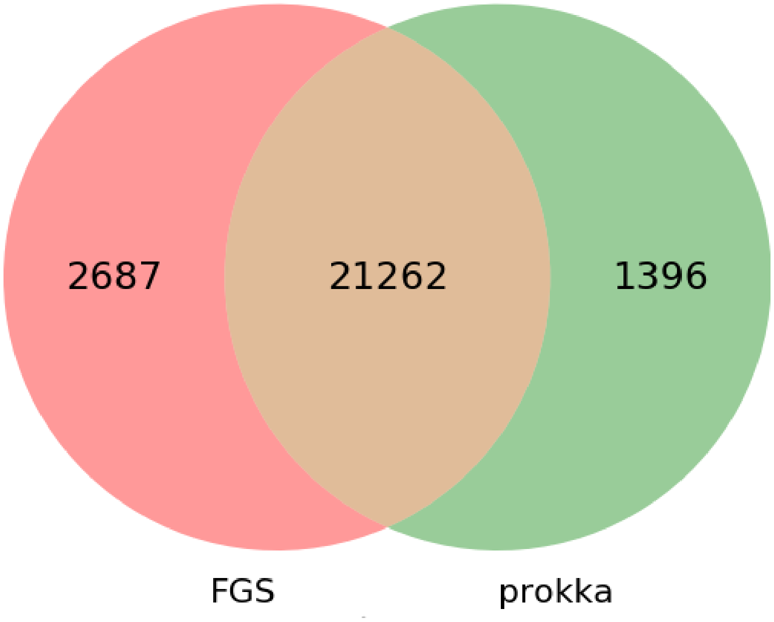


Figure S3: Venn diagram summarizing the overlap between the rescued ORFs that were missed by FragGeneScan and prokka.

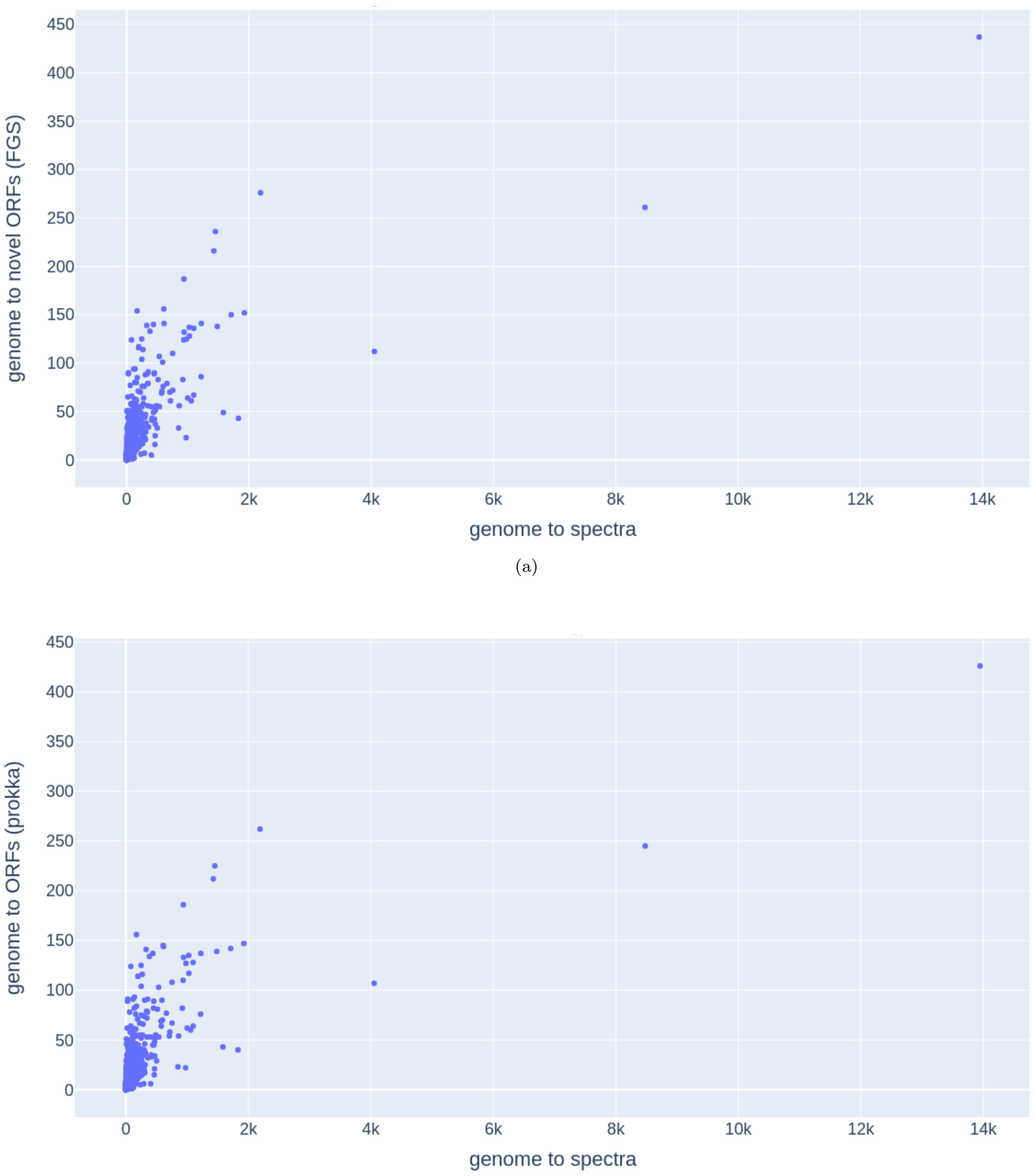


(a)

(b)

Figure S4: Scatter plots showing the relationship between the expression levels of the different genomes at the protein level with the total number of rescued ORFs. (A) rescued ORFs missed by FragGeneScan; (B) rescued ORFs missed by prokka.
